# Long-range massively parallel reporter assay reveals rules of distal enhancer-promoter interactions

**DOI:** 10.1101/2025.04.21.649048

**Authors:** Yawei Wu, Jie Li, Emma L Bartley-Dier, Cassidy Pitts, Barak A Cohen

## Abstract

Long-range activation is an essential property of enhancers, yet the features determining long-range enhancer activities have not been systematically investigated due to a lack of high-throughput methods to measure long-range enhancer activities efficiently. To address this gap, we present a long-range massively parallel reporter assay (long-range MPRA), an assay allowing the measurement of hundreds of enhancers at multiple distances from a genome-integrated promoter. The long-range MPRA assay features two independent landing pads, allowing modular control over the genome-integrated promoter and enhancer libraries. We showcased the capability of long-range MPRA by testing over 300 K562 enhancers, as well as a set of enhancer combinations, at distances up to 100 kb. We found that enhancers’ long-range activities are primarily determined by their intrinsic strength, with strong enhancers retaining more activity over long distances, while weak enhancers rapidly lose activity. Additionally, we found that GATA1-bound enhancers are more resistant to distance-dependent loss of activity, suggesting that TF binding also modulates long-range function. Finally, testing long-range enhancer activities with three different promoters (*HBE*, *HBG* and *GAPDH*) revealed that long-range enhancer-promoter interactions rely on not only enhancer properties but also promoter responsiveness.

## Introduction

Enhancers are cis-regulatory elements that coordinate gene expression programs through interactions with their target promoters. A remarkable feature of enhancers is their ability to regulate genes over long genomic distances, ranging from several kilobases to megabases^1–3^. These long-range enhancer-promoter (E-P) interactions are prevalent in the mammalian genome, with two-thirds of the ENCODE-annotated human enhancers being more than 10 kb away from the nearest genes^4^. Many long-range enhancers play important roles in development. Disruption of existing E-P interactions, or induction of ectopic interactions, can both lead to developmental dysfunctions^1,5,6^.

Despite the prevalence and importance of long-range E-P interactions, it remains unclear what rules govern long-range E-P communication. 3D genome organization, constructed by cohesin-mediated, CTCF-anchored structural loops, is thought to facilitate E-P interactions by bringing E-Ps into physical proximity. However, a series of studies have challenged this view by showing that depletion of cohesin or CTCF has little impact on global E-P interactions or gene expression^7–10^. Local disruptions of the 3D genome have also shown mixed effects: while some enhancers rely on preformed structural loops^11,12^, many enhancers still function at a distance on their own^13–16^. This varying reliance on pre-established proximity suggests that enhancers have sequence-encoded intrinsic ability to drive long-range interactions^17^. Understanding what determines an enhancer’s intrinsic long-range capability is critical for predicting enhancer activity across different genomic contexts, as well as determining how enhancer activity might change in response to alterations of enhancer sequence or 3D genome structure.

Different approaches have been used to study features of long-range enhancers. Chromatin contact capture techniques^18–21^ and CRISPR-based enhancer perturbations^22–24^ helped identify a catalog of functional E-P connections. Genome-wide computational analyses of those E-P connections have revealed the enrichment of certain transcription factor (TF) motifs in long-range enhancers^25,26^. Concurrently, detailed characterizations of individual long-range enhancers, conducted in their endogenous loci^27–30^ or in synthetic contexts^31–34^, have confirmed the role of specific TFs, as well as highlighting the importance of enhancer strength. Additionally, studies on enhancer clusters also revealed the importance of cooperativity among multiple enhancers^31,35^. These studies have generated valuable hypotheses about the potential features of long-range enhancers. However, these hypotheses derive from either correlative studies, which lack direct causality, or isolated examples, which may not be generalizable. Moreover, the roles of features such as enhancer strength, TF motif content, and enhancer cooperativity are not mutually exclusive, and are often confounded with each other and by other variables, such as different genomic contexts, making it difficult to dissect the contribution of each individual feature. A systematic approach is therefore needed to directly test the role of these features in a controlled genomic context.

Beyond enhancer features, another important question is whether long-range E-P communication requires specific promoter properties. Two recent studies have tested E-P compatibility on plasmid-based reporter assays and showed that most enhancers work well with most promoters with little selectivity^36,37^. However, because these studies positioned enhancers adjacent to their targeted promoters, it remains unknown whether the same lack of E-P selectivity applies over larger genomic distances. Addressing this question requires an assay capable of measuring long-range enhancer activities with the flexibility to test different promoters in a controlled context.

To address these questions, we developed a long-range massively parallel reporter assay (long-range MPRA). Long-range MPRA is a genome-integrated reporter assay that allows modular control over enhancers and promoters, and the distance between these regulatory elements. Thus, long-range MPRA allows the measurement of hundreds of enhancers, at multiple E-P distances, in parallel. To demonstrate the assay’s capability to explore rules of long-range E-P interactions, we measured over 300 enhancers from K562 cells at 0, 10 and 100 kb from a *HBG* promoter-driven reporter gene. We found that at 10 kb, enhancer strength was the primary determinant of enhancer activity, but TF binding also played a role, with GATA1-bound enhancers being more resistant to distance-caused activity loss. At 100 kb, the strength requirement became more stringent, with only the strongest enhancers remaining active. Additionally, by testing a set of enhancer combinations, we found that synergistic interactions among multiple enhancers could confer robust activation at distances up to 100 kb, yet their activity still adhered to the strength-dependent rule observed for single enhancers. Finally, comparing long-range enhancer activities across three promoters (*HBE*, *HBG* and *GAPDH*) revealed that promoters differ in their responsiveness to long-range enhancers: *HBE* promoter was robustly activated by enhancers integrated at 0 kb, but showed much weaker response at 10 kb; *HBG* was very responsive at both distances; and *GAPDH*, a strong housekeeping promoter, showed minimal induction at either distance. Altogether, our results demonstrate that long-range enhancer function is determined by a combination of enhancer strength, TF binding, and promoter responsiveness.

## Results

### Design of long-range MPRA

Our goal was to develop a reporter assay allowing efficient testing of enhancer libraries at various distances from any promoter of interest. We previously established “landing pad” MPRA for integrating and measuring enhancers in a pre-built loxP cassette in the genome (i.e. the landing pad) through recombinase-mediated cassette exchange (RMCE)^38–40^. Inspired by that, we designed long-range MPRA, featuring a dual landing pad system where enhancers and promoters can be integrated independently at two separated landing pads (**Fig. 1a**). We achieved orthogonal integrations of enhancers and promoters through two distinct recombination systems: Flp-FRT^41^ for enhancer integration and Cre-loxP^42^ for promoter integration (**Fig. 1a**). Each landing pad also includes unique fluorescence and counterselection markers to facilitate precise and efficient selection of cells that have undergone the desired recombinations (**Fig. 1a**). This dual landing pad design allows flexible control of the enhancer sequence, the promoter sequence, and the distance between the two.

**Fig. 1.**
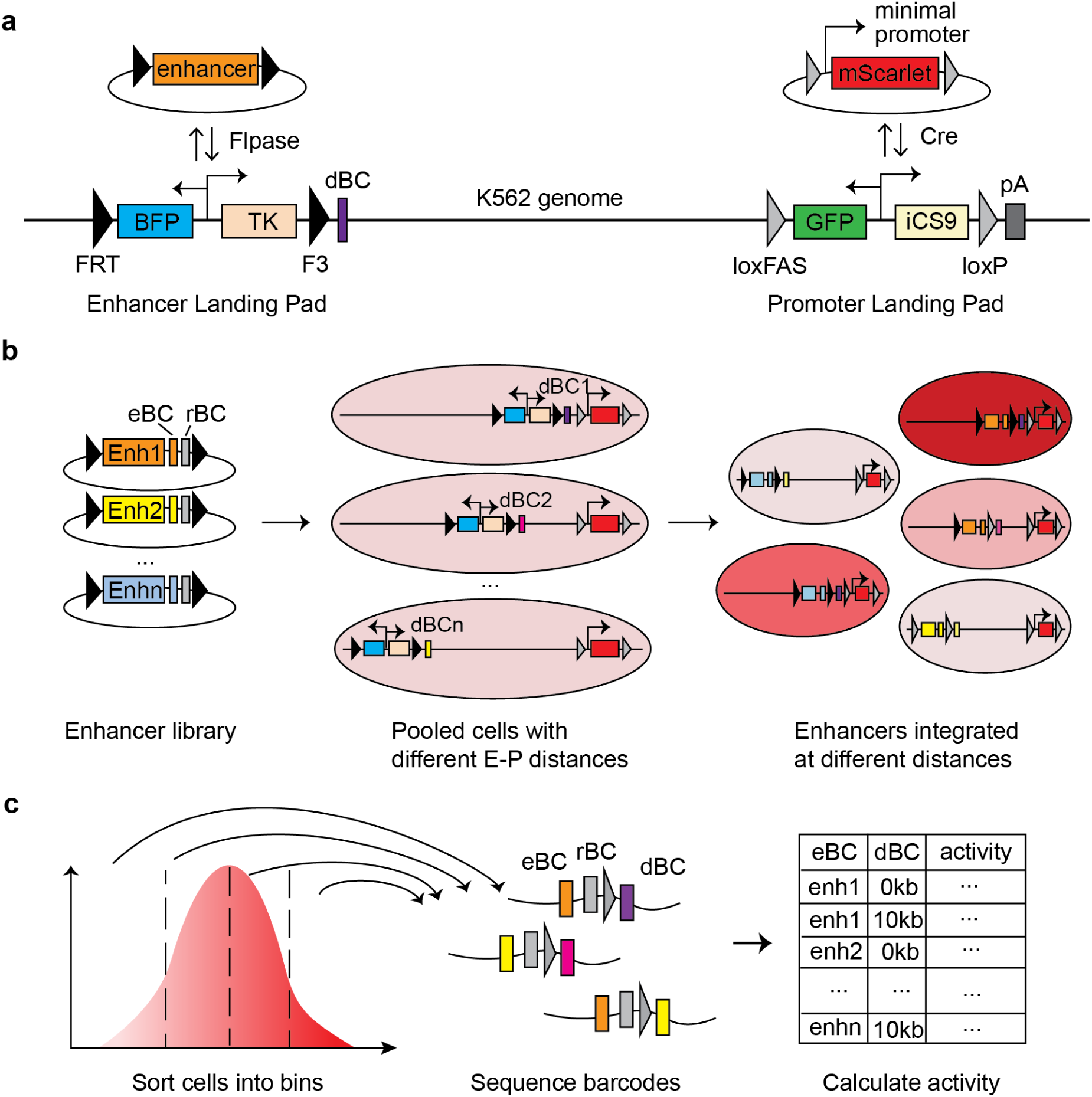
Design of long-range MPRA. **a**, Design of the enhancer landing pad and the promoter landing pad. The enhancer landing pad contains a dual promoter driving expression of an mTagBFP fluorescent protein and a thymidine kinase (TK) gene for counterselection, flanked by FRT and F3 sites for RMCE. The promoter landing pad expresses the mEmerald fluorescent protein and iCaspase9 for counterselection and is flanked by loxFAS and loxP sites for orthogonal RMCE. **b-c,** Workflow of long-range MPRA. eBC: enhancer barcode; rBC: random barcode; dBC: distance barcode. A barcoded enhancer library is integrated into pooled long-range landing pad cells through Flpase-facilitated recombination. After counterselection with Ganciclovir, cells with successful integration are enriched. Cells are sorted into different bins based on mScarlet fluorescence level. Genomic DNA is extracted from each bin; barcodes are amplified and sequenced. Reporter gene expression of each enhancer at each distance is calculated as the weighted average of fluorescence levels across bins.

To multiplex the measurement of many enhancers at multiple E-P distances, we use an enhancer barcode (eBC) to identify each enhancer in the enhancer library, a distance barcode (dBC) to specify the genomic distance of the enhancer landing pad relative to the promoter, and a random barcode (rBC) that contains 16 Ns to track individual integration events (**Fig. 1b**). After integrating a minimal-promoter driven fluorescence reporter at the promoter landing pad, and integrating a library of barcoded enhancers into pooled cells with enhancer landing pads at different distances (**Fig. 1a,b**), we can use sort-seq^23,43–45^ to connect the barcode information at the enhancer landing pad with the expression output from the promoter landing pad (**Fig. 1c**). Specifically, by sorting cells into different bins based on fluorescent reporter expression and then sequencing the eBC-rBC-dBC barcode regions in each bin, we can determine the distribution of the barcodes across the bins. The reporter gene expression for each enhancer at each E-P distance can therefore be calculated as the weighted average fluorescence intensity across the bins (**Fig. 1c**).

We constructed our long-range MPRA system in K562 cells on chromosome 18 (chr18:58014732), located within an intergenic region with minimal active enhancers, promoters, or CTCF sites to avoid potential interference from the surrounding environment (**Extended Data Fig. 1a**). We engineered the two landing pads by two sequential CRISPR knock-ins: first knocking in the promoter landing pad and then adding the enhancer landing pad at 3, 10, 20, 50 and 100 kb upstream of the promoter landing pad. Cell lines with 0 kb E-P distance were made separately by co-integrating the enhancer and promoter landing pad with a single conjugated CRISPR donor sequence (**Methods**). With this dual landing pad system, the promoter sequence of interest is first recombined into the promoter landing pad to generate reporter cell lines, which are subsequently used to test enhancer libraries at varying distances.

### At 10 kb, both enhancer strength and GATA1 binding affect long-range enhancer activity

Using long-range MPRA, we first sought to explore what type of enhancers can work over long distances. Specifically, we aimed to test enhancers of varying strengths and TF binding profiles to determine whether long-range activation needs specific TFs, or if enhancer strength alone confers function. We focused on GATA1, a master erythroid TF known to bind many long-range enhancers in K562 cells, including the well-known β-globin locus control region (LCR)^27,28,46^. GATA1 interacts with LDB1, which is reported to have self-association ability and can mediate extreme long E-P interactions^28,29,47–50^. Additionally, GATA1 binding and GATA1 motifs are enriched in distal enhancers compared to proximal ones in K562 cells^25^. Together, these pieces of evidence suggest a potential role of GATA1 binding in long-range enhancer function.

To test the contributions of enhancer strength and GATA1 binding, we constructed a library containing 180 GATA1-bound enhancers and 130 non-GATA1-bound enhancers, selected based on ENCODE K562 H3K27ac and GATA1 ChIP-seq peaks (**Fig. 2a**, **Methods**)^4^. To evaluate the influence of enhancer strength, enhancers in both groups were chosen to span a range of activities in the ENCODE K562 MPRA assay (**Extended Data Fig. 2a**, **Methods**)^4,51^. Additionally, we included 40 GATA1-bound enhancers with mutated GATA1 motifs, 20 dinucleotide-shuffled sequences as negative controls, and several known enhancers tested in previous MPRA assays as positive controls (**Fig. 2a**)^36,51,52^. In total, our library contains 383 unique 249 bp sequences. We ordered them as 350nt oligos to include constant sequences for cloning, a 10 bp unique enhancer barcode (eBC) to identify each library member, and a 16 bp random barcode (rBC) to track unique integration events.

**Fig. 2.**
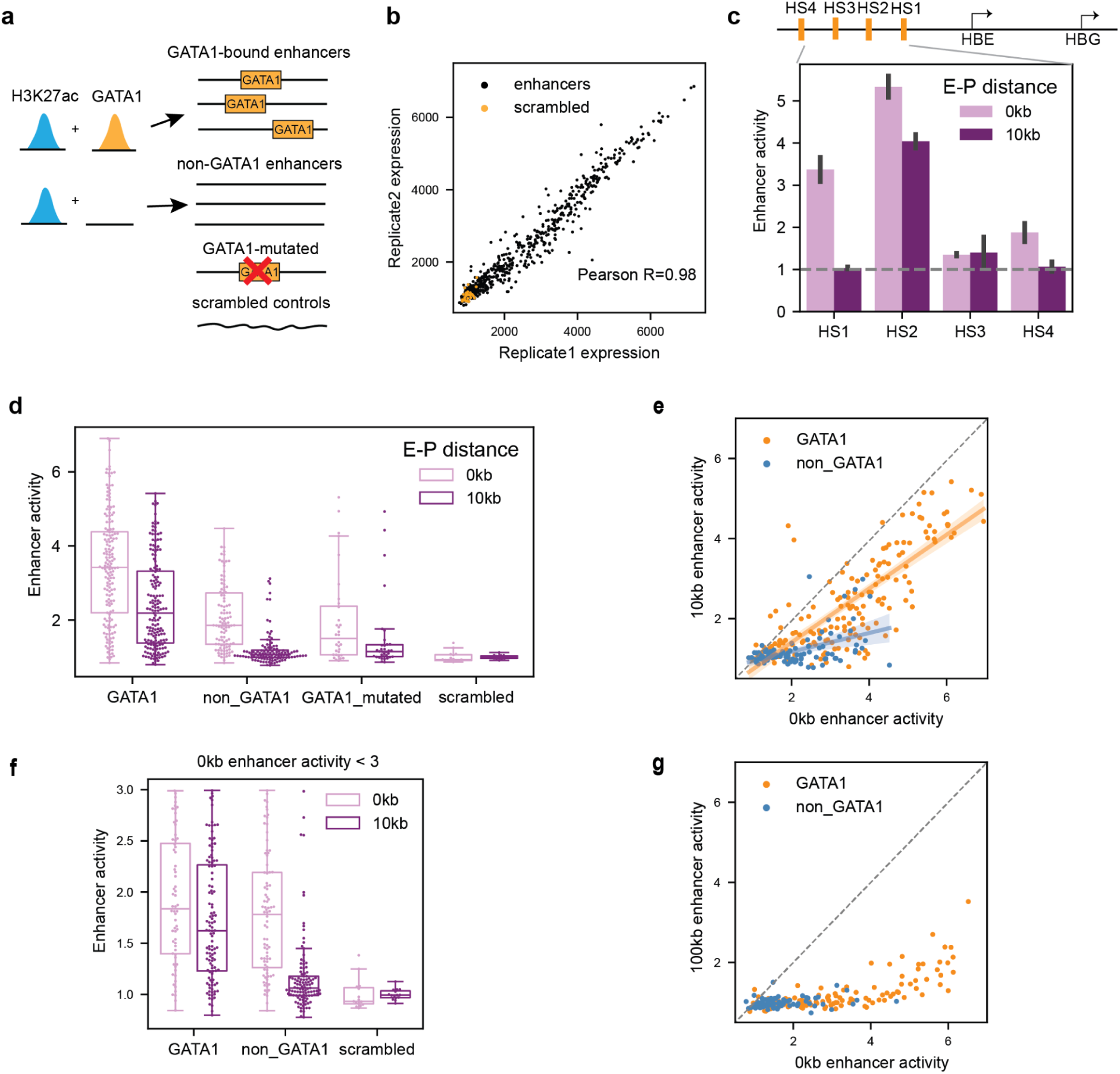
At 10 kb, both strength and TF binding affect long-range enhancer activity. **a**, Design of the GATA1/non-GATA1 enhancer library. We selected GATA1/non-GATA1-bound enhancers by overlapping K562 H3K27ac ChIP-seq peaks with GATA1 peaks. We also picked 40 GATA1-bound enhancers and mutated all the GATA1 motifs to assess the effect of GATA1 binding. We made 20 dinucleotide shuffled sequences as negative controls. **b,** Correlation of long-range MPRA measurements between two biological replicates, calculated for each individual enhancer integrated at either 0 kb or 10 kb. Axes represent the measured reporter expression, which was calculated as weighted average mScarlet fluorescence intensity across sorted bins. Scrambled controls are highlighted in orange. **c,** 0 kb and 10 kb enhancer activity of HS1,2,3 and 4 of the β-globin LCR. Bars show the average enhancer activity of two replicates. Enhancer activity was calculated as reporter expression fold change over scrambled controls. 0 kb and 10 kb are colored light purple and dark purple, respectively. Error bars are 95% CI (n=2). The dashed grey line shows y=1. **d,** Box plot of 0 kb and 10 kb enhancer activity of different enhancer groups. Boxes represent the median, IQR, and whiskers (±1.5 IQR). **e,** Correlation of 10 kb versus 0 kb activity. The dashed grey line is y=x. GATA1 and non-GATA1 enhancers are colored orange and blue, respectively. **f,** 0 kb and 10 kb enhancer activity of a subset of enhancers with 0 kb activity smaller than 3. Boxes are the same as (d). **g,** Correlation of 100 kb versus 0 kb activity. The dashed grey line is y=x. GATA1 and non-GATA1 enhancers are colored orange and blue, respectively.

We assayed this GATA1/non-GATA1 enhancer library at 0 kb and 10 kb upstream of a γ-globin gene (*HBG*) promoter-driven mScarlet reporter (**Methods**). The *HBG* promoter is an erythroid promoter regulated by β-globin LCR, a well-known long-range enhancer in K562 cells, making it a biologically relevant reporter for testing long-range enhancer function. We used 0 kb activity as a proxy for enhancer strength, and 10 kb activity to evaluate long-range performance. From long-range MPRA, we successfully recovered 98% of library members. By analyzing the number of unique rBCs per eBC-dBC pair, we estimated that each enhancer was integrated a median of 63 times at each distance. To ensure robust measurements, we only kept enhancers with >10 integrations at both 0 kb and 10 kb, resulting in 86% of the library used for downstream analyses. We calculated the reporter expression for each enhancer at each distance as the weighted average fluorescence intensity across sorted bins.

Long-range MPRA measurements of the GATA1/non-GATA1 enhancer library were highly reproducible between two biological replicates (Pearson *R* = 0.98), and spanned a broad expression range compared to scrambled controls (**Fig. 2b**). For downstream analyses, we quantified enhancer activity as fold change relative to scrambled controls, representing each enhancer’s direct contribution to expression above baseline promoter activity. This assay also successfully detected known enhancers and long-range enhancers. Four positive controls – DNase hypersensitive sites (HSs) within the β-globin LCR known to regulate *HBG* promoter – all activated *HBG* reporter at 0 kb, and their rankings were consistent with previous episomal MPRA studies^36,51^ (**Fig. 2c**). Also, long-range MPRA identified HS2 as the only fragment exhibiting robust long-range activity at 10 kb, which agrees with the previous finding that HS2 is essential for LCR-mediated long-range activation (**Fig. 2c**)^53^.

To validate long-range MPRA measurements, we selected 20 enhancers from the library, individually integrated them at 0 kb and 10 kb, and measured reporter expression using flow cytometry. We confirmed that individually measured enhancer activities correlated well with long-range MPRA measurements (Pearson *R* = 0.91 at 0 kb, 0.90 at 10 kb, **Extended Data Fig. 3a**). In addition, we performed RT-qPCR to assess 0 kb and 10 kb activity of 7 enhancers and confirmed that long-range MPRA readouts reflect true RNA-level changes (Pearson *R* = 0.78 at 0 kb, 0.86 at 10 kb, **Extended Data Fig. 3b**).

After confirming the accuracy and reproducibility of the measurements, we next used this dataset to investigate how enhancer strength and GATA1 binding contribute to long-range enhancer activity. Firstly, by comparing GATA1– and non-GATA1-bound enhancers, we found that GATA1-bound enhancers exhibited higher activities at both 0 kb and 10 kb (**Fig. 2d**). While enhancer activity decreased at 10 kb for both groups, non-GATA1-bound enhancers were more severely affected, with the majority losing all activity, whereas many GATA1-bound enhancers remained active (**Fig. 2d**). Mutating GATA1 motifs in GATA1-bound enhancers resulted in significant reduction in activity at both 0 kb and 10 kb, bringing their levels down to the levels of non-GATA1 enhancers, suggesting that GATA1 binding is necessary for both enhancer strength and long-range performance (**Fig. 2d**).

We next sought to determine whether GATA1-bound enhancers’ superior long-range activity is purely a consequence of their higher strength, or if GATA1 binding confers additional contributions. When plotting enhancers’ 10 kb activity against their 0 kb strength, we found a strong correlation (Pearson *R* = 0.82, **Fig. 2e**), indicating that enhancer strength is a strong predictor of 10 kb enhancer activity. Indeed, using ANOVA, we found that strength alone could explain 66% of the variance observed in 10 kb activity (*P* = 2.44 x 10^-64^), suggesting that much of the higher 10 kb activity for GATA1-bound enhancers can be attributed to their stronger baseline strength.

Despite the overall correlation between 10 kb activity and strength, when we examined GATA1– and non-GATA1-bound enhancers separately, we found that on average GATA1-bound enhancers retained 67% of their initial activity from 0 kb, whereas non-GATA1-bound enhancers retained only 20% (compare slopes in **Fig. 2e**). Moreover, when GATA1 binding status was incorporated into the ANOVA analysis as a slope-modifying factor, the explained variance increased from 66% to 71%, with the additional contribution of GATA1 binding being statistically significant (*P =* 1.07 x 10^-9^), confirming that GATA1 contributes beyond simply enhancing strength. GATA1’s effect was particularly evident in a subset of weak enhancers (**Fig. 2f**), where GATA1– and non-GATA1 enhancers exhibited similar 0 kb strength (*P* = 0.16, *t-*test), but GATA1-bound enhancers showed significantly higher 10 kb activities (*P* = 9.12 x 10^-15^, *t-*test). Together, these findings reveal that GATA1’s role in long-range enhancer function is multifaceted–it contributes to enhancer strength and also improves an enhancer’s resilience to increased distances.

### At 100 kb, enhancers require higher strength to remain active

We observed a strong correlation between enhancer strength and their long-range activity at 10 kb, but it is unclear if this relationship holds at even greater distances or if additional features are required to overcome a larger genomic barrier. To test that, we measured the GATA1/non-GATA1 enhancer library at 0 kb and 100 kb away from the *HBG* promoter using long-range MPRA. We chose 100 kb as a representative of extreme long distance because it lies in the long tail of E-P distance distribution–only ∼10% of K562 enhancers are located more than 100 kb away from the nearest promoters^4^.

At 100 kb, most enhancers exhibited little to no activity, with only a handful of the strongest GATA1 enhancers remaining active, although at low levels (**Fig. 2g**). We observed a non-linear correlation between 100 kb activity and enhancer strength (Spearman *R* = 0.72, **Fig. 2g**), suggesting that (1) enhancer strength continues to be a key determinant of long-range activation, and (2) increasing distance disproportionately affects weak enhancers compared to strong ones. Unlike at 10 kb, where weak enhancers (especially GATA1-bound ones) retained a range of activities (**Fig. 2e,f**), at 100 kb, almost all weak enhancers became inactive regardless of their GATA1 binding status (**Fig. 2g**). This indicates that at extreme distances, the strength requirement becomes more stringent, and TF binding alone is insufficient to override this requirement.

Notably, the enhancer library we tested already included the top 100 GATA1 enhancers and the top 70 non-GATA1 enhancers with the strongest strength in ENCODE lentiMPRA assays^4,51^ (**Methods**), yet at 100 kb only a handful were active. Although it is possible that synthetic enhancers can achieve stronger strength^54,55^, our result still suggested that a single genomic enhancer, with its length limited to only 249 bp, may not have enough strength to confer robust activation over extreme distances. Indeed, many well-characterized examples of long-range activation involve locus control regions that contain multiple copies of enhancers^31,35,53^. Therefore, we sought to move from single enhancers to enhancer combinations, to test if cooperativity among multiple enhancers can confer better long-range performance.

### Synergistic interactions among multiple enhancers confer robust long-range activity through increasing enhancer strength

In the genome, enhancers often appear in clusters, where multiple enhancers are positioned adjacent to each other. Many of these enhancer clusters activate gene expression over large genomic distances^35,56,57^. From our results with the GATA1/non-GATA1 enhancer library, we observed a correlation between single enhancers’ strength and their long-range performance. Here we sought to determine whether cooperativity among multiple enhancers can boost long-range performance, and whether this increased long-range activity still adheres to the same strength-dependent pattern observed for single enhancers.

To address this question, we focused on the β-globin LCR, a well-characterized enhancer cluster containing four GATA1-bound DNase hypersensitive sites (HSs). Each HS has been shown to exhibit short-range activity in previously published plasmid MPRAs, as well as in our long-range MPRA (**Fig. 2c**). Perturbation of any HS at LCR’s endogenous locus affects downstream globin gene expression^22,23^. To systematically test how combinations of these four HSs influence long-range activity, we amplified 1.5 kb fragments centered around DNase-seq peaks of HS1, HS2, HS3 and HS4 from the genome and constructed a library containing various combinations of the four HSs: (1) single HSs: HS1, HS2, HS3 and HS4; (2) pairs: HS12, HS13, HS14, HS23, HS24 and HS34; (3) triplets: HS123, HS124, HS134 and HS234; (4) the full LCR: HS1234. We also included a no enhancer control in the library to indicate basal promoter expression. To obtain a more continuous view of how enhancer activity responds to increased distances, we tested this library at 0, 3,10, 20, 50 and 100 kb upstream of the *HBG* promoter using long-range MPRA **(Fig. 3a**).

**Fig. 3.**
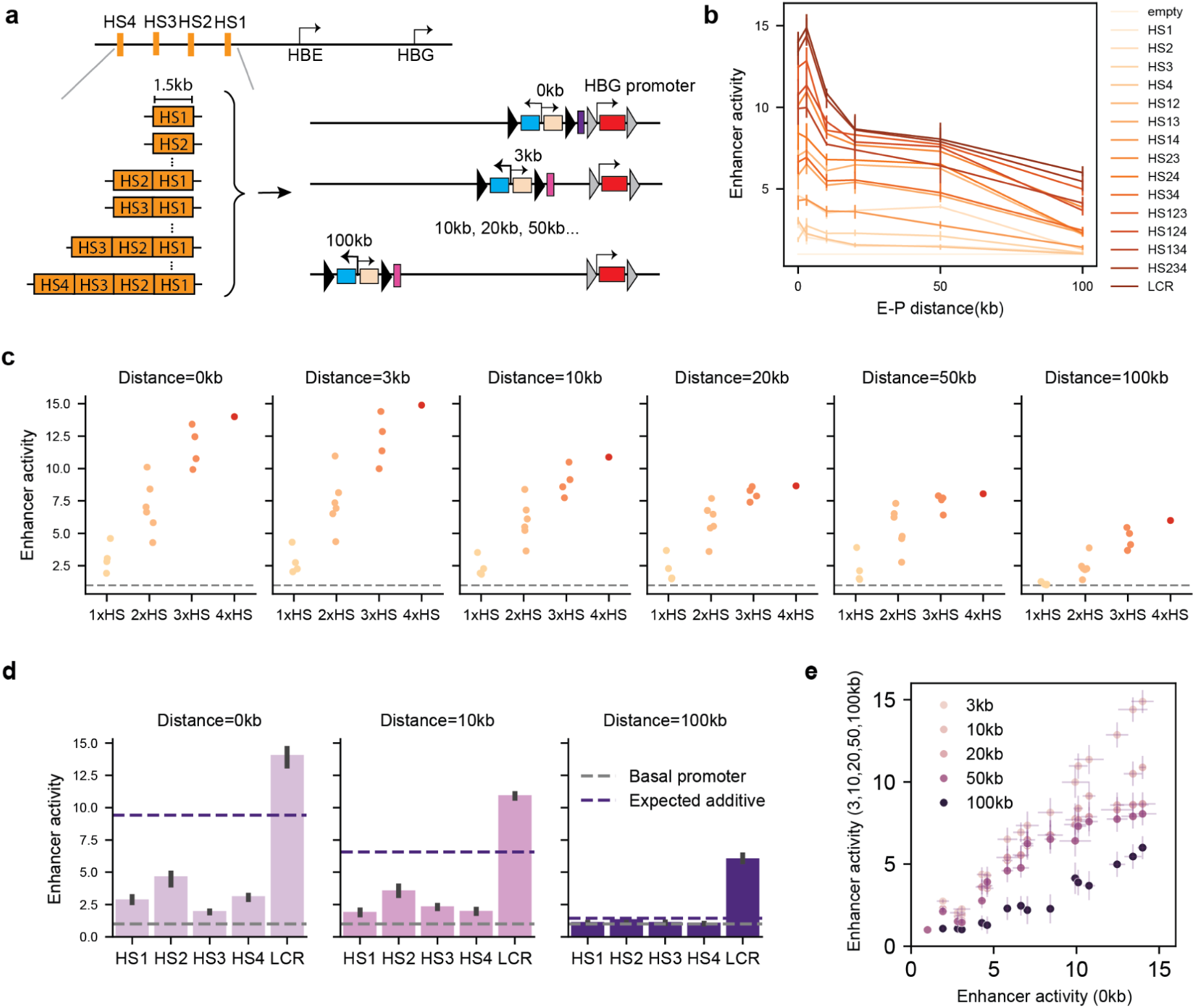
Synergistic interactions among multiple enhancers confer robust long-range activity through increasing enhancer strength. **a**, Design of the LCR HS combination library. We PCR amplified 1.5 kb fragments of HS1, 2, 3, and 4 from the β-globin locus, and cloned 16 combinations of these fragments. We integrated the library into mixed 0, 3, 10, 20, 50, and 100 kb landing pad cells and measured their activities using long-range MPRA. **b,** Enhancer activities of each library member across the 6 distances. Error bars are 95% CI (n=3). Lines are colored by enhancer identity. **c,** Enhancer activities of fragments containing different numbers of HSs. The X-axis shows the number of HSs (from 1x HS to 4x HS). The Y-axis shows enhancer activity at the corresponding distance. The dashed grey line is y=1. **d,** Bar plot of 0, 10, and 100 kb activities of single HSs and full LCR. Error bars are 95% CI (n=3). Dashed purple lines indicate expected LCR activity if the four HSs act additively. Dashed grey lines are y=1. **e,** Correlation of 3, 10, 20, 50, 100 kb versus 0 kb activity. Dot darkness represents distance, with darker purple indicating longer distance. Error bars are 95% CI (n=3).

As the E-P distance increased, enhancer activity decreased in a nonlinear manner for all elements, including individual HSs and their combinations (**Fig. 3b**). Overall, combining more HSs led to higher 0 kb strength and higher long-range activity across all distances (**Fig. 3c**). The four HSs cooperated synergistically at both short and long distances, with their combined activity exceeding the sum of individual activities (**Fig. 3d**). This synergy was most dramatic at 100 kb, where single HSs had little to no detectable activity, whereas the full LCR still retained robust activity (**Fig. 3d**).

Similar to our observations with single enhancers, we found a strong correlation between long-range activities of the LCR HSs and their initial strength at 0 kb (**Fig. 3e**). Distance scaled enhancer activity in a strength-dependent manner: at a short distance such as 3 kb, the relationship between enhancer activity and strength was nearly linear; at 100 kb, weak enhancers, including single HSs and weak combinations such as HS14, were more severely affected than strong enhancers (**Fig. 3e**). This nonlinear pattern at 100 kb is similar to what we observed with single enhancers (**Fig. 2e**). The difference is that the LCR HS combination library extended the range of enhancer strength, such that the strongest single enhancers from the GATA1/non-GATA1 library now fell within the weaker range of the LCR HS combination library. These findings reinforce that enhancer strength is a key determinant of long-range enhancer performance, even when multiple enhancers are combined. However, we should note that all four HSs we tested bind GATA1 (**Extended Data Fig. 4a**), and thus likely represent a scenario where enhancers with similar TF binding profiles are combined. It is possible that certain combinations of enhancers that recruit distinct TFs/cofactors could enhance long-range activity through alternative mechanisms beyond merely increasing enhancer strength.

### Test of enhancers against different promoters reveals promoter-specific long-range responses

Next, we asked if an enhancer’s long-range performance is affected by what promoter it interacts with. Previous studies have suggested that at short-range there is little E-P selectivity^36,37^. It is not clear if this lack of selectivity also extends to long-range E-P interactions. In the experiment where we tested a library of GATA1/non-GATA1 enhancers with *HBG* promoter, we found that GATA1-bound enhancers outperformed non-GATA1 enhancers at both short– and long-range. We asked whether this finding is specific to *HBG*, which is an erythroid-specific promoter known to be regulated by GATA1-bound enhancers, or if it also applies to other promoters. To address this question, we measured the same GATA1/non-GATA1 enhancer library with two additional promoters: *HBE*, another erythroid promoter also regulated by LCR, and *GAPDH*, a housekeeping promoter that is not regulated by any GATA1-bound enhancers (**Extended Data Fig. 4**). The three promoters exhibited different basal activities when integrated into the promoter landing pad alone, with *GAPDH* being the strongest and *HBE* being the weakest (**Fig. 4a**).

**Fig. 4.**
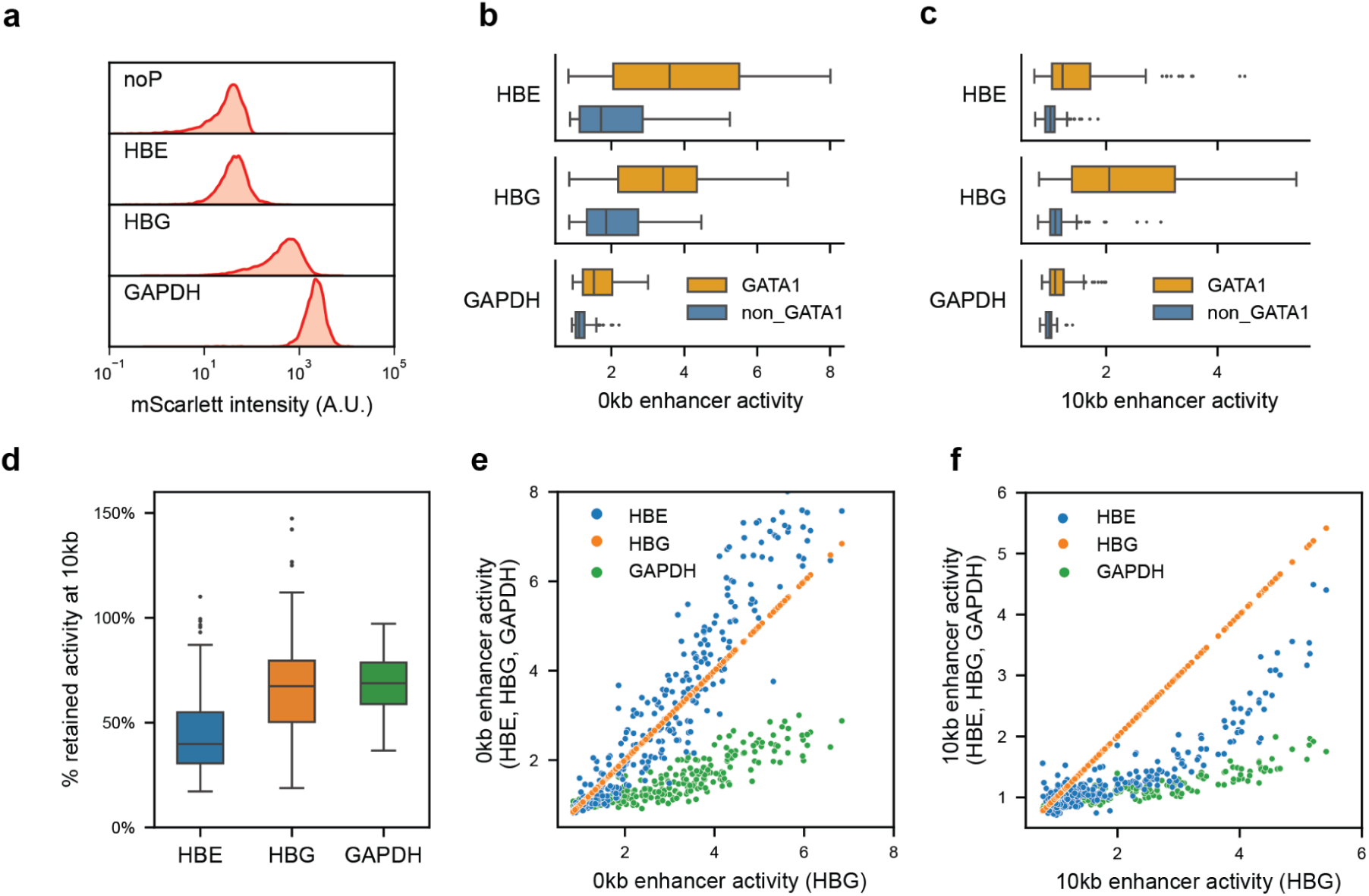
Test of enhancers against different promoters reveals promoter-specific long-range responses. **a**, mScarlet fluorescence intensity of cells integrated with no promoter (noP), *HBE*, *HBG* and *GAPDH* promoter. We obtained fluorescence measurements from Sony SY3200 sorter and plotted distribution of PE-A signal at log scale. **b-c,** Box plot of 0 kb (b) or 10 kb (c) enhancer activities for three different promoters. Orange indicates GATA1 enhancers and blue indicates non-GATA1 enhancers. Boxes represent the median, IQR, and whiskers (±1.5 IQR). **d,** Percentage of retained activity at 10 kb versus 0 kb, calculated as 10 kb enhancer activity divided by 0 kb activity. Boxes represent the median, IQR, and whiskers (±1.5 IQR). **e-f,** Correlation of 0 kb (e) or 10 kb (f) enhancer activity with *HBE* (blue), *HBG* (orange) and *GAPDH* (green) promoter versus with *HBG* promoter.

When enhancers were integrated immediately upstream of the promoter, we found that the overall induction of promoter activity inversely correlates with the promoter’s basal activity (**Fig. 4b**). Specifically, *GAPDH*, the strongest promoter, showed the smallest induction (< 2-fold), while *HBE*, the weakest promoter, was most responsive to enhancers, showing an induction of up to 8-fold. This trend, where stronger promoters are less responsive to further activation, is consistent with previous findings from our lab and other studies^36,37,39^.

Surprisingly, promoters’ responses at 10 kb did not follow the same pattern as observed at 0 kb. While *GAPDH* continued to be the least induced promoter, likely due to its already small induction at 0 kb, the order of the other two promoters switched. *HBE* promoter, despite being highly inducible at 0 kb, showed much weaker responses to 10 kb enhancers, while *HBG* became the most responsive promoter at 10 kb (**Fig. 4c**). Comparing the percentage of retained enhancer activity when moved from 0 kb to 10 kb, enhancers paired with *HBG* retained on average 68% of the activity, whereas those paired with *HBE* retained only 45% (**Fig. 4d**).

Although the three promoters showed different overall responsiveness to enhancers at short/long distances, GATA1-bound enhancers continuously exhibited stronger short-range and long-range activities than non-GATA1 enhancers (**Fig. 4b,c**). This suggested that GATA1 enhancers’ higher strength and superior long-range performance are not specific to GATA1-regulated promoters such as *HBG* and *HBE*, but can extend to housekeeping promoters as well. Moreover, when directly comparing enhancer activities across promoters, we found that the relative ranking of enhancer activity remained remarkably consistent at both 0 kb and 10 kb (**Fig. 4e,f**), suggesting that these three promoters, despite differing in overall responsiveness, do not exhibit strong preferences for specific enhancers. However, we observed that *HBE*’s decreased responsiveness at 10 kb was more pronounced for weak enhancers (**Fig. 4f**). This nonlinear pattern suggests that strong enhancers might partially compensate for the low long-range responsiveness of the *HBE* promoter. Altogether, our results showed that long-range activation is a combined result of both enhancer capability and promoter responsiveness.

## Discussion

We present long-range MPRA as a new addition to the enhancer assay toolbox, enabling high-throughput measurements of enhancer activities over long genomic distances. Short-range enhancer assays have greatly improved our understanding of sequence determinants of enhancer strength, as well as enhancer-promoter compatibility. However, it remains a question how much of the rules learned at short-range can extend to long-range E-P interactions. Long-range MPRA fills this gap by providing a high-throughput platform to test rules governing long-range E-P interactions in a controlled genomic context.

Using long-range MPRA, we examined how TF binding, enhancer strength, enhancer cooperativity, and promoter type affect an enhancer’s long-range capability. Among these factors, our results revealed enhancer strength as one of the strongest determinants of whether an enhancer can effectively activate a promoter over long distances. Across all tested conditions–from 3 kb to 100 kb, with single enhancers or enhancer combinations, and against different promoters–long-range activity was consistently correlated with basal enhancer strength. This correlation is near linear at moderate distances such as 10 kb, whereas at greater distances, such as 100 kb, the strength requirement becomes more stringent, with enhancers below a certain threshold showing no detectable activity. This strength-dependent long-range capability was also demonstrated by Thomas et al., who tested nine enhancers of different strength at distances of 1.5 kb, 25 kb, and 75 kb^31^. Our findings expand their observations to over 300 enhancers, as well as a handful of enhancer combinations, indicating that this strength-dependent long-range activation is a common phenomenon. At the molecular level, a strong enhancer’s ability to maintain long-range activity likely comes from its recruitment of more transcriptionally active proteins, including TFs, Pol II transcriptional machinery, and the associated cofactors. Many of these transcription-associated proteins contain intrinsically disordered domains (IDRs) that mediate multivalent interactions^58^. These interactions might facilitate long-range E-P communication by forming large protein complexes that bridge E-Ps, increasing E-P encounter frequency, or promoting cohesin loading/stalling to stabilize cohesin-mediated loops^33,59^. Future studies coupling long-range MPRA with chromatin contact capture and live cell imaging techniques will help better understand the molecular mechanisms behind this strength-conferred long-range activity.

Although enhancer strength is a key factor, it does not fully explain long-range activity. For example, at 10 kb, weak GATA1-bound enhancers have significantly higher long-range activity than equally weak enhancers lacking GATA1 binding, suggesting that the identity of recruited proteins also affects long-range performance. GATA1 is known to form complexes with TAL1, LMO2, and LDB1^28^, with LDB1 previously shown to mediate long-range E-P interactions in various cell types. Structurally, LDB1 contains a LIM interaction domain, which can interact with other LIM domain proteins, and a dimerization domain that self-interacts^28,47^. It is likely that the recruited LDB1 at GATA1-bound enhancers promotes E-P looping through its LIM domain and dimerization domains. Similar LDB1-mediated long-range E-P interactions have been found in the olfactory bulb, where LDB1 interacts with LHX2 to mediate trans-chromosomal interactions between olfactory receptor genes and their enhancer hubs^50^. Additionally, LDB1 and LHX9 are likely involved in the recently identified REX element (range-extender element) that facilitates long-range E-P interactions in the limb bud^32^. Interestingly, while LHX motifs in the REX element are required specifically for long-range activity, but not for short-range^32^, GATA1 motifs are required for both short– and long-range as shown in our results. It is potentially because GATA1-LDB1 and LHX-LDB1 complexes differ in their transcriptional activities in different cell types.

Our experiments with three different promoters suggest that long-range E-P interactions depend not only on enhancer properties but also on promoter responsiveness. It is previously reported that promoter strength affects how strongly it can be activated by enhancers on plasmid-based assays^36,37,39^. Therefore, it is not surprising that the strong housekeeping promoter *GAPDH* is poorly activated by enhancers at both short and long distances, due to its already high basal activity. However, *HBE* promoter, although robustly induced at 0 kb, responded poorly at 10 kb, especially to weak enhancers. This suggests that long-range activation requires additional promoter properties. Similar to enhancers, the number (strength) and identity of recruited proteins at a promoter may affect how easily it can interact with proteins recruited by distant enhancers. One possibility is that a promoter needs to recruit enough protein mass to form a hub, so that it can be more easily captured by enhancers. This may explain why the combination of a very weak promoter (*HBE*) and a weak enhancer fails to create critical protein mass required for 10 kb activation. Alternatively, *HBG* promoter might contain motifs that bind essential looping factors that do not bind *HBE* promoter. Investigating the determinants of long-range promoter responsiveness would require systematic testing of many promoters with varying strength and different sequence features at long distances, which can be done by testing barcoded promoter libraries at the promoter landing pad using long-range MPRA.

Here we demonstrated the capability of long-range MPRA to explore sequence-encoded features in enhancers and promoters. In the future, with some innovations, long-range MPRA can be adapted to studying many other players involved in long-range gene regulation. For instance, by creating libraries that include auxiliary elements, such as insulators that block E-P interactions, or “facilitators” (e.g., REX element) that promote them, we can systematically dissect the roles of these elements in long-range activation. Moreover, by comparing long-range enhancer activities across various genomic contexts: active versus repressive, within versus across TAD boundaries, we can test whether long-range E-P communication follows distinct rules in different contexts. Additionally, coupling long-range MPRA with protein perturbations, such as targeted recruitment or depletion of specific TFs/cofactors, will allow us to directly dissect the roles of trans-regulatory factors in long-range E-P interactions.

## Methods

### Cell culture

Wild-type K562 erythroleukemia cells were obtained from the Genome Engineering & Stem Cell Center at Washington University in St. Louis, and were maintained in Iscove’s Modified Dulbecco′s Medium (IMDM) + 10% FBS + 1% nonessential amino acids (Gibco #11140050) + 1% penicillin/streptomycin. Cells were passaged every three days to maintain a density of 0.2-1 million cells per ml and were regularly tested for mycoplasma following the PCR protocol described in Young et al^60^.

### Long-range landing pad cell line construction

The long-range landing pad cell lines contain two independent integration systems(landing pads): loxFAS-mEmerald-iCS9-loxP for promoter integration and FRT-TagBFP-TK-F3 for enhancer integration(**Fig. 1a**). We constructed these two landing pads through two sequential rounds of CRISPR-mediated knock-ins.

First, we integrated the promoter landing pad (loxFAS-mEmerald-iCS9-loxP-WPRE-polyA) at chr18:58014732. We designed sgRNAs for knock-ins using Benchling and cloned them into the pX330 plasmid (Addgene #42330) via the Bbs1 sites. The knock-in donor plasmid (pYWD26) contains the promoter landing pad flanked by 750 bp homology arms. We transfected 1.2 million wild-type K562 cells with 1 μg cloned pX330 plasmid and 3 μg donor plasmid using the Neon Transfection System 100 μL kit (Invitrogen #MPK10096). One week after transfection, we sorted mEmerald+ cells into single-cell clones and confirmed correct heterozygous knock-ins using PCR.

With the promoter landing pad successfully integrated, we added the enhancer landing pad at 3, 10, 20, 50, and 100 kb upstream of the promoter landing pad using CRISPR. The enhancer landing pad contains FRT-TagBFP-TK-F3, followed by a unique 12 bp barcode for each of the distances, and a constant primer binding site for barcode PCR. For each E-P distance, we transfected 1.2 million K562 cells already containing the promoter landing pad with 1 μg cloned pX330 plasmid and 3 μg corresponding donor plasmid using the Neon Transfection System 100 μl kit (Invitrogen #MPK10096). After transfection, TagBFP+/mEmerald+ cells were sorted into single clones and screened by PCR to confirm heterozygous integration.

0 kb landing pad cells were individually constructed by co-integrating the enhancer and the promoter landing pad with a single conjugated CRISPR donor sequence (pYWD28).

To confirm cis configuration of the two landing pads, for 3, 10, and 20 kb, we used one primer in the enhancer landing pad and one primer in the promoter landing pad to amplify across the interval. The presence of a long PCR product indicates integration on the same chromosome. For 50 and 100 kb distances, we employed digital droplet PCR (ddPCR) to check physical linkage of the two landing pads, following BioRad’s QX200 protocol for variant phasing. More specifically, we extracted high molecular weight genomic DNA from candidate clones using the Monarch HWM DNA extraction kit, and used HEX– and FAM-labeled probes to amplify regions in the enhancer and promoter landing pads. Clones with correct cis integration will show enriched double-positive droplets in ddPCR.

DNA sequences of all sgRNAs, primers, probes and plasmids mentioned above can be found in the **Supplementary Table 1**.

### Promoter cloning and integration into promoter landing pad

We cloned a promoter transfer backbone (pYWD30) that contains loxFAS-Age1-Nhe1-mScarlet-loxP. We obtained promoter sequences of *HBG*, *HBE,* and *GAPDH* by extracting upstream 20 bp and downstream 195 bp sequences of the RefSeq annotated TSS (see **Supplementary Table 1** for promoter sequences). We ordered these promoter sequences as gBlocks from IDT and cloned them individually between loxFAS and mScarlet in the Age1&Nhe1 digested pYWD30 using NEBuilder HIFI assembly (NEB, E2621).

To integrate a specific reporter gene into the promoter landing pad, we transfected long-range landing pad cells with 2ug pCMV-Cre (Addgene, #123133) and 6ug promoter transfer plasmid. 3 days after transfection, we started treating cells with 1nM AP20187 (MCE #HY-13992, diluted in DMSO) for three days to kill cells still expressing iCS9, and then sorted TagBFP+/mEmerald-cells into single clones.

### GATA1/non-GATA1 enhancer library design and cloning

We intersected ENCODE K562 DNase-seq peaks (ENCFF185XRG) with H3K27ac ChIP-seq peaks (ENCFF544LXB) to obtain a candidate list of enhancers^4^. To remove promoter-like elements, we filtered the candidate list based on ENCODE cCRE registry for K562 cells (ENCFF210CAN). Next, we intersected candidate enhancers with K562 GATA1 ChIP-seq peaks (ENCFF657CTC) to divide enhancers into GATA1-bound and non-GATA1-bound groups. The GATA1-bound enhancers were further scanned for instances of GATA1 motifs using FIMO^61^ (version 5.4.1, p-value cutoff=0.025) to ensure they are directly bound by GATA1.

Among these candidate GATA1-bound enhancers, we selected 180 sequences to span a range of potential enhancer activities: including 100 enhancers with the strongest enhancer activity in ENCODE K562 MPRA data (ENCSR382BVV)^4,51^, and another 80 enhancers sampled evenly across the range of MPRA activity. Additionally, we included 13 GATA1-bound enhancers within +-100 kb regions of several important erythroid genes (GATA1, HBB, LMO2) as positive controls; we also included the mutated version of 40 GATA1-bound enhancers where all the GATA1 motifs were mutated. From non-GATA1-bound enhancers, we selected 130 sequences in total, including 70 DNase peaks with the strongest MPRA activities and 60 other enhancers spanning a range of activities. We also included 20 dinucleotide-shuffled oligos as negative controls.

All together, the GATA1/non-GATA1 enhancer library contains 383 library members. We selected 249 bp centered around the middle of the H3K27ac peak and assigned one unique 10 bp enhancer barcode (eBC) to label each individual library member. We ordered the library as a 350nt oligo pool from IDT. Each oligo is constructed according to the following template: 5’ primer annealing sequence/Age1/enhancer/BamH1/Sph1/eBC/Xho1/3’ primer annealing sequence. Sequences and barcode information of all library members can be found in **Supplementary Table 2**.

To clone the oligo library into a plasmid, we first cloned an enhancer transfer backbone (pYWD16) that contains FRT-Age1-Xho1-F3. We PCR amplified the oligo library using Q5 High-Fidelity 2X Master Mix (NEB, M0942) with primers lr_P14: CCTATTCTCTAGAAAGTATAGGAACTTCACCGGT and lr_P15: GATCCTCCTCTCTGCCACACCC. We performed PCR in 8 separate 50 μl reactions using the following conditions: 98 °C for 30 s; 4 cycles of 98 °C for 15 s, 69 °C for 15 s, and 72 °C for 45 s. We gel purified the amplified library and then cloned the purified library into Age1&Xho1 digested pYWD16 backbone using NEBuilder HiFi assembly (NEB, E2621). We digested the cloned library with Sbf1 (NEB, 3642S) and then inserted a 16 bp random barcode using HiFi assembly following the manufacturer’s protocol for ssDNA cloning.The ssDNA oligo used for barcode insertion was: lr_P73: CTCGAGCTCGGGTGTGGCAGAGAGGAGGATCCNNNNNNNNNNNNNNNNCCTGCAGGGAAGCGCTAGCGAAGTTCCTATAC.

### LCR library design and cloning

HS1, HS2, HS3, and HS4 were amplified from genomic DNA using primers targeting +/-750 bp from the center of the four DNase peaks at the LCR locus. 14 plasmids containing different combinations of the four HSs were individually cloned into pYWD16 backbone using NEBuilder HiFi DNA Assembly (NEB, E2621), including (1) 4 plasmids containing only one HS: HS1, HS2, HS3 and HS4; (2) 6 plasmids containing two HSs: HS1&2, HS1&3, HS1&4, HS2&3, HS2&4 and HS3&4; (3) 3 plasmids containing three HSs: HS1&2&3, HS1&2&4 and HS2&3&4; (4) 1 plasmid containing all four HSs: HS1&2&3&4. We also cloned one plasmid where the center 380 bp of the HS2 fragment is deleted, and one plasmid containing only the center 380 bp HS2 sequence. Additionally, the empty backbone (pYWD16) that doesn’t contain any enhancer is used as the negative control.

After all 18 plasmids were cloned, a 10 bp enhancer barcode (eBC) was added to individual plasmids by inserting a 10 bp random sequence downstream of the enhancer using Q5 site-directed mutagenesis kit (NEB, E0554). We selected one clone with a distinct barcode for each of the 17 library members. All barcoded plasmids were manually pooled together in equal ratios and then digested with Sbf1 (NEB, E3642S); a 16 bp random barcode (rBC) is then cloned to the Sbf1 site using NEBuilder HiFi DNA Assembly.

Sequences and barcode information of all library members can be found in **Supplementary Table 2**.

### Enhancer library integration and sort-seq measurement

To test enhancer libraries at multiple E-P distances, we first mixed long-range landing pad cell lines of selected E-P distances in equal ratios and transfected the pooled cells with the enhancer library and Flpase recombinase using the Neon Transfection System 100 μl kit. We performed two or three replicates for each enhancer library. For each replicate, we performed 12 transfections, each with 1.2 million cells and a mix of 2 μg pCAG-Flpe (Addgene, #13787) and 6 μg enhancer library plasmid. We also did one control transfection with Flpase and a transfer plasmid with no enhancer to indicate the activity of the basal promoter. On Day 3 after transfection, we started to treat cells with 2uM Ganciclovir (sigma, 82410-32-0, diluted in water) for ten days to kill cells without enhancers integrated.

On Day 14 we sorted cells on a Sony SY3200 sorter at the Siteman Cancer Center Flow Cytometry Core. We set an ON/OFF threshold for mScarlet as the 10th percentile of the basal promoter expression. The lowest bin was set as cells below 75% of the control. The ON region was further split into 3 bins at a 40%:30%:20% ratio (**Extended Data Fig. 5**). For each replicate, we collected 1.5 million cells in each of the four bins. We noted the median fluorescence intensity and the percentage of cells in each bin and used those numbers to calculate enhancer activity.

Genomic DNA was extracted from cells collected from 4 bins for each replicate using QIAamp DNA mini kit (Qiagen, #51304). For each DNA sample, we performed 8 separate 50 μl reactions with lr_P3: GCATGCGAATGGCCTCGTAG and lr_P5: CACAGCCAGACCAACATTGCC using the following conditions: 98 °C for 30 s; 20 cycles of 98 °C for 15 s, 69 °C for 15 s, and 72 °C for 45 s. After PCR, we pooled eight PCRs for each sample, cleaned up DNA using 1.2x SPRIselect beads (Beckman Coulter, B23317), and quantified DNA concentration using Qubit (Invitrogen, Q33231). We added sequencing adapters and indices with two additional rounds of PCRs: for PCR2, we took 4ng PCR1 product as template and performed 8 cycles of PCR with primers lr_P4: CTTTCCCTACACGACGCTCTTCCGATCTGCATGCGAATGGCCTCGTAG and lr_P6: GTGACTGGAGTTCAGACGTGTGCTCTTCCGATCTCACAGCCAGACCAACATTGCC. For PCR3, we took 2ul PCR2 product as template and amplified another 8 cycles using primers IDT_10_i5: AATGATACGGCGACCACCGAGATCTACAC(10 bp index)ACACTCTTTCCCTACACGACGCTC and IDT_10_i7: CAAGCAGAAGACGGCATACGAGAT(10 bp index)GTGACTGGAGTTCAGACGTG. After PCR3, we cleaned up DNA using 1.2x SPRIselect beads and quantified DNA concentration using Qubit (Invitrogen, Q33231). We sequenced pooled PCR3 products on the Illumina NextSeq platform or the Element Biosciences AVITI platform, using paired-end reads of 98 bp from Read 1 and 50 bp from Read 2.

### Barcode sequencing data analysis

We parsed the enhancer barcode (eBC), distance barcode (dBC), and random barcode (rBC) from each read in the fastq files and counted the abundance of each eBC-dBC-rBC combination. Because rBC is highly complex, each eBC-dBC-rBC combination represents a unique integration event. We filtered out eBC-dBC-rBC combinations with <= 20 reads to remove under-sequenced integrations or potential sequencing errors, and then only kept eBC-dBC pairs with >= 10 unique rBCs to make sure every enhancer at every distance was measured from at least 10 independent integration events.

After filtering, we collapsed all rBCs associated with certain eBC-dBC pairs. We normalized the read counts by the total reads in each bin and then renormalized each barcode across bins to get barcode distribution probability. We used the distribution probability and the median mScarlet fluorescence of each bin to calculate the weighted average mScarlet level of each eBC-dBC pair. To compare enhancer activities across different distances, we normalized the calculated mScarlet level by the mScarlet level of the basal promoter with no-enhancer control or with scrambled controls. When comparing GATA1/non-GATA1 library performance at 0&10 kb versus 0&100 kb, we applied an additional normalization to align the mean 0 kb value between the two datasets, ensuring that the shared 0 kb enhancer measurements spanned the same range. All calculated enhancer activity values used in the paper are provided in **Supplementary Table 4**.

### Individual enhancer validation with flow cytometry and RT-qPCR

We selected 20 enhancers from the GATA1/non-GATA1 enhancer library that exhibited a range of 0 kb and 10 kb activities in long-range MPRA (see **Supplementary Table 3** for enhancer identities). We ordered these enhancers as eblocks from IDT and individually cloned them into the pYWD16 backbone. We integrated those enhancers, as well as pYWD16 as a no-enhancer control, at 0 kb/10 kb upstream of the *HBG* promoter by transfecting 1.2 million 0 kb/10 kb landing pad cells with 2 μg Flpase and 6 μg enhancer transfer plasmids. We performed two separate transfections for each enhancer as biological replicates.

After treating cells with Ganciclovir for 7 days followed by an additional 4 days of culture, we measured tagBFP and mScarlet fluorescence levels using Beckman Coulter’s Cytoflex S flow cytometer. Flow cytometry data were analyzed using CytExpert software. Reporter gene expression was quantified as the mean PE-A value of BFP negative population and enhancer activity was calculated by normalizing fluorescence levels by the no-enhancer control.

In addition to flow cytometry measurements, we extracted RNA from a subset of cells integrated with 7 enhancers and the no-enhancer control using the Monarch Total RNA Miniprep Kit (NEB #T2010). We measured mScarlet and housekeeping gene HPRT using the Luna Universal One-Step RT-qPCR kit (NEB #E3005) with 100 ng total RNA per reaction, two reactions per replicate (mScarlet primers: lr_P69: CCCGTAATGCAGAAGAAGACA, lr_P70: GCGCAGGGCCATCTTAAT; HPRT primers: lr_P71: GTATTCATTATAGTCAAGGGCATATCC, lr_P72: AGATGGTCAAGGTCGCAAG). Enhancer activity was calculated using the ΔΔCt method, normalized to the no-enhancer control.

### Epigenetic feature analysis

We downloaded the foldchange-over-control bigWig files for K562 H3K27ac, H3K4m3, POLR2A, GATA1, TAL1, EP300, ATF1 and CTCF ChIP-seq from ENCODE^4^. We used pyBigWig to calculate the mean histone/TF binding signal at the endogenous locations of all the enhancers in our library, as well as a list of non-enhancer regions as negative controls. For each ChIP-seq signal, we standardized the values and then plotted the average signal for GATA1-bound enhancers, non-GATA1-bound enhancers and negative controls respectively in a heatmap using seaborn.heatmap.

### Statistics and data visualization

All the statistics tests were performed in Python using Numpy^62^, Scipy^63^ and Statsmodels^64^. Correlation coefficients were calculated using scipy.stats.pearsonr and scipy.stats.spearmanr. Linear regression for enhancers’ 0 kb and 10 kb activities were done using scipy.stats.linregress. *T-*tests were performed using scipy.stats.ttest_ind or scipy.stats.ttest_rel.

To determine the contribution of strength and GATA1 binding to long-range enhancer activity, we performed ANOVA using statsmodels.api.stats.anova_lm. We first fitted a baseline model: 10 kb activity ∼ 0 kb activity + residual, to assess how much variance in 10 kb activity can be explained by 0 kb strength alone. Then we fitted a new model with GATA1 binding status incorporated as a slope modifier: 10 kb activity ∼ 0 kb activity + 0 kb activity:C(GATA1_binding) + residual, to determine how much additional variance can be explained by GATA1 binding.

All the figures were generated in Python using Matplotlib^65^ and Seaborn^66^.

## Data availability

Raw sequencing data and barcode counts have been deposited to the NCBI Gene Expression Omnibus (GEO) under accession GSE294367. All processed data used in this study are available at GitHub (<u>yaweiwu233/long-range-MPRA</u>). K562 DNase-seq, ChIP-seq, RNA-seq tracks and rE2G predicted E-P connections were downloaded from ENCODE^4^. K562 HiC data was obtained from ENCODE (ENCFF080DPJ)^4^ and visualized using UCSC genome browser.

## Code availability

The code used to process all barcode sequencing data, and to generate all the figures in this study is available at GitHub (yaweiwu233/long-range-MPRA).

## Acknowledgements

We thank members of the Cohen Lab and Dustin Baldrige Lab for helpful discussions and critical feedback on the manuscript. We are grateful to Jessica Hoisington-Lopez and MariaLynn Crosby at the DNA Sequencing Innovation Lab at the Center for Genome Sciences & Systems Biology for assistance with DNA sequencing; Brian Koebbe and Eric Martin for computing cluster support; staff at the Alvin J. Siteman Cancer Center at Washington University School of Medicine and Barnes-Jewish Hospital for assistance with cell sorting; Shiming Chen and Susan Penrose with the Molecular Genetics Service Core for their technical support with digital droplet PCR. This work was supported by National Institutes of Health grants R01GM092910, R01GM140711, and R01HG012304 to B.A.C.

## Author contributions

Y.W. and B.A.C. conceived and designed the project. Y.W., J.L., E.L.B.D. and C.P. conducted the experiments. Y.W. performed the analyses. Y.W. and B.A.C. wrote the manuscript.

## Extended Data Figures

**Extended Data Fig. 1:**
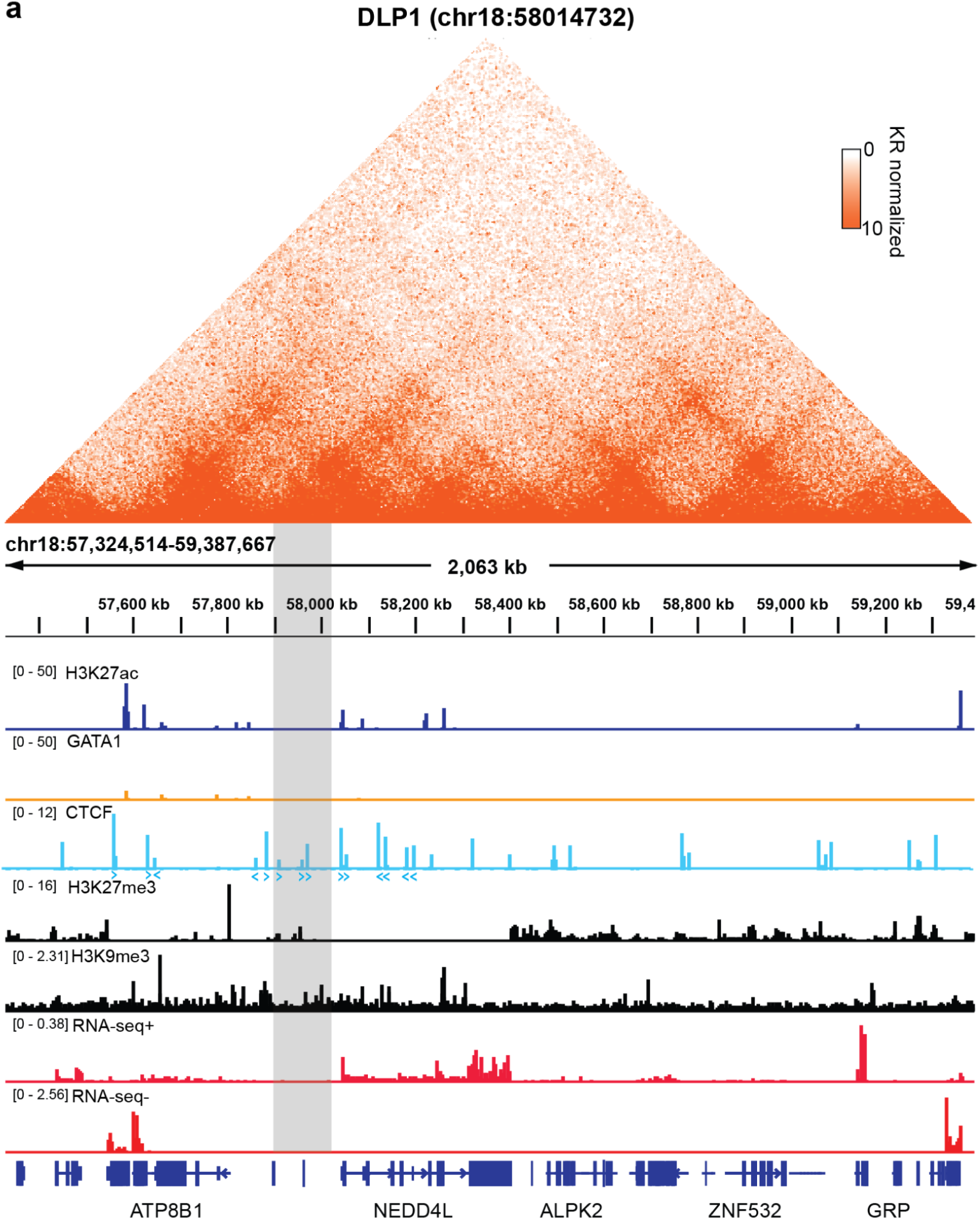
Genomic context of landing pads in K562 cells. **a**, The promoter landing pad is integrated at chr18:58014732. Enhancer landing pads are integrated at 0, 3, 10, 20, 50 and 100 kb upstream of the promoter landing pad. The region containing landing pads is indicated by the shaded area. The top shows Knight-Ruiz (KR) normalized HiC contact map for K562 cells at 5 kb resolution taken from ENCODE. Tracks show: ChIP-seq for H3K27ac, GATA1, CTCF, H3K27me3 and H3K9me3, total RNA-seq data for plus/minus strand and RefSeq gene annotations. CTCF motifs near landing pad locations are indicated by arrows.

**Extended Data Fig. 2:**
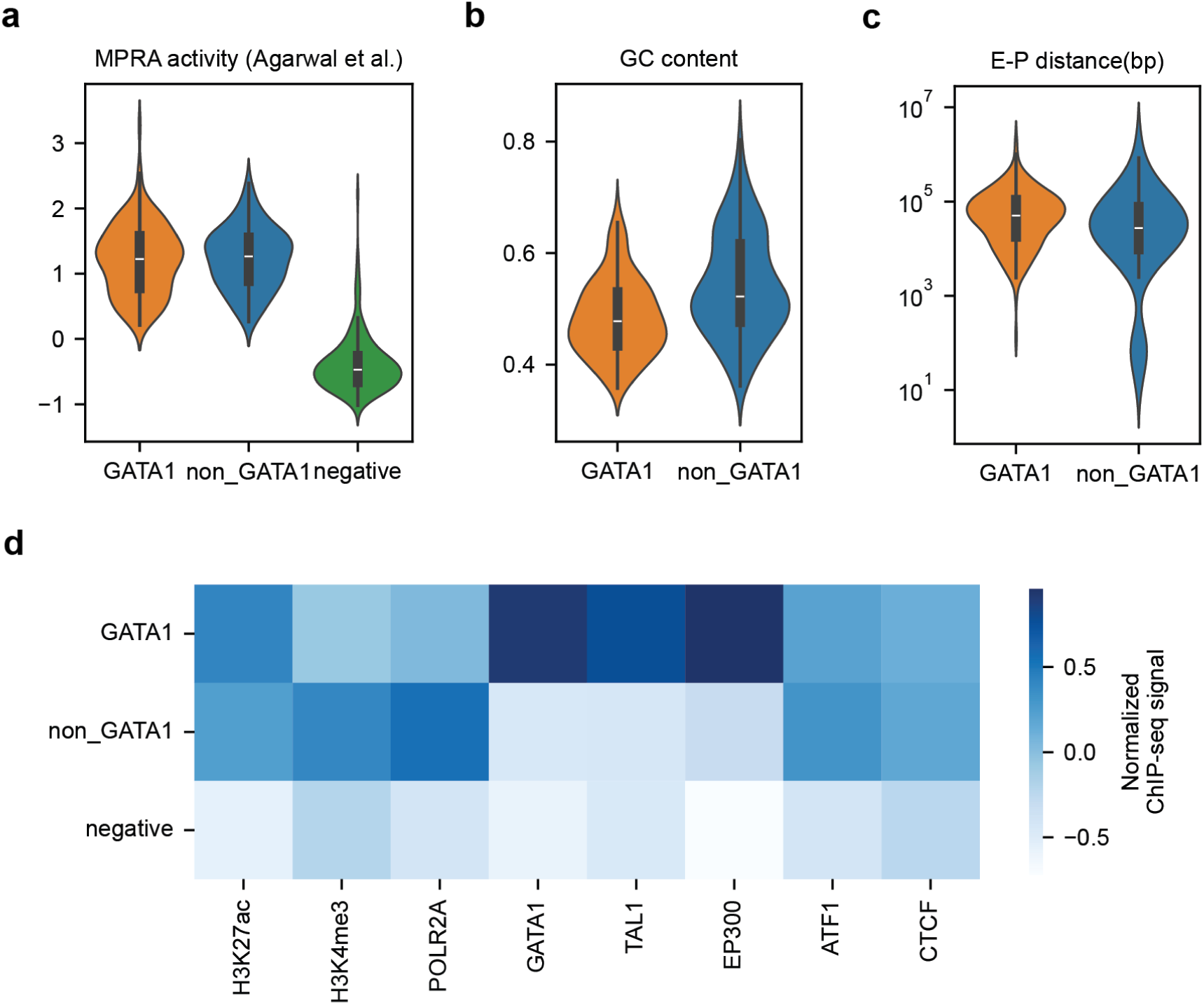
Sequence and epigenetic features of GATA1-bound versus non-GATA1-bound enhancers in the library. **a**, The violin plot shows MPRA activities, measured as log2-transformed RNA/DNA, for GATA1– and non-GATA1-bound enhancers in our GATA1/non-GATA1 library. We downloaded K562 MPRA data from ENCODE. Negative controls are control genomic sequences used in the ENCODE MPRA library. **b,** The violin plot shows GC content of GATA1– and non-GATA1-bound enhancers. **c,** The violin plot shows the distribution of distance to the nearest active promoter for GATA1– and non-GATA1-bound enhancers. Y-axis is the log10-transformed distance (bp). **d,**The heatmap shows the mean normalized ChIP-seq signal of several histone marks, cofactors and TFs.

**Extended Data Fig. 3:**
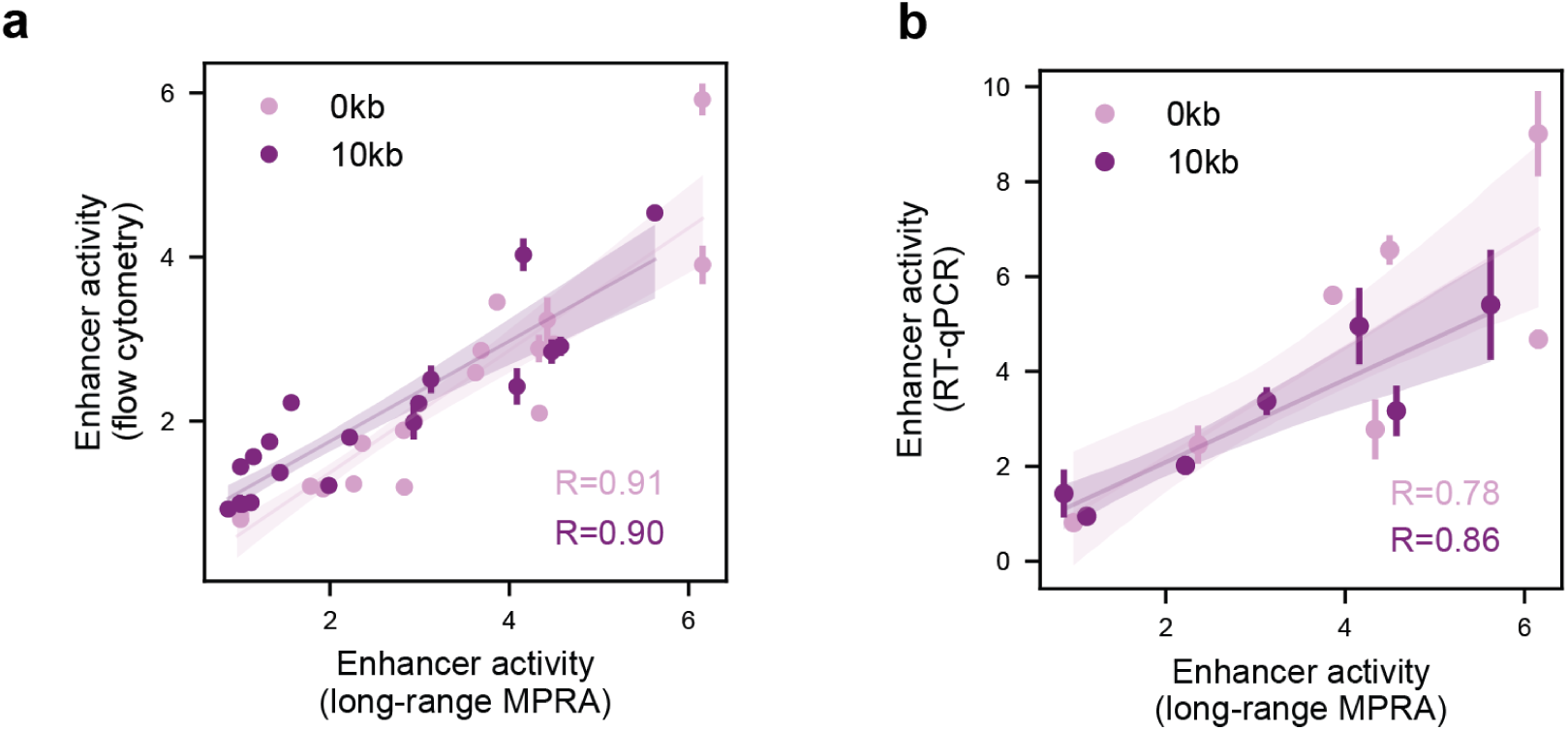
Individual enhancer validation with flow cytometry and RT-qPCR. **a**, The scatter plot shows the correlation between enhancer activity measured by long-range MPRA (X-axis) and by flow cytometry (Y-axis). Each dot represents an enhancer integrated at either 0 kb (light purple) or 10 kb (dark purple). The error bars represent 95% confidence interval (n=2). The two lines show the linear regression fits, with the translucent bands around the regression line showing the 95% confidence interval of the regression, estimated by seaborn.regplot. **b**, Same as in (a), but the Y-axis shows the enhancer activity measured by qRT-PCR.

**Extended Data Fig. 4:**
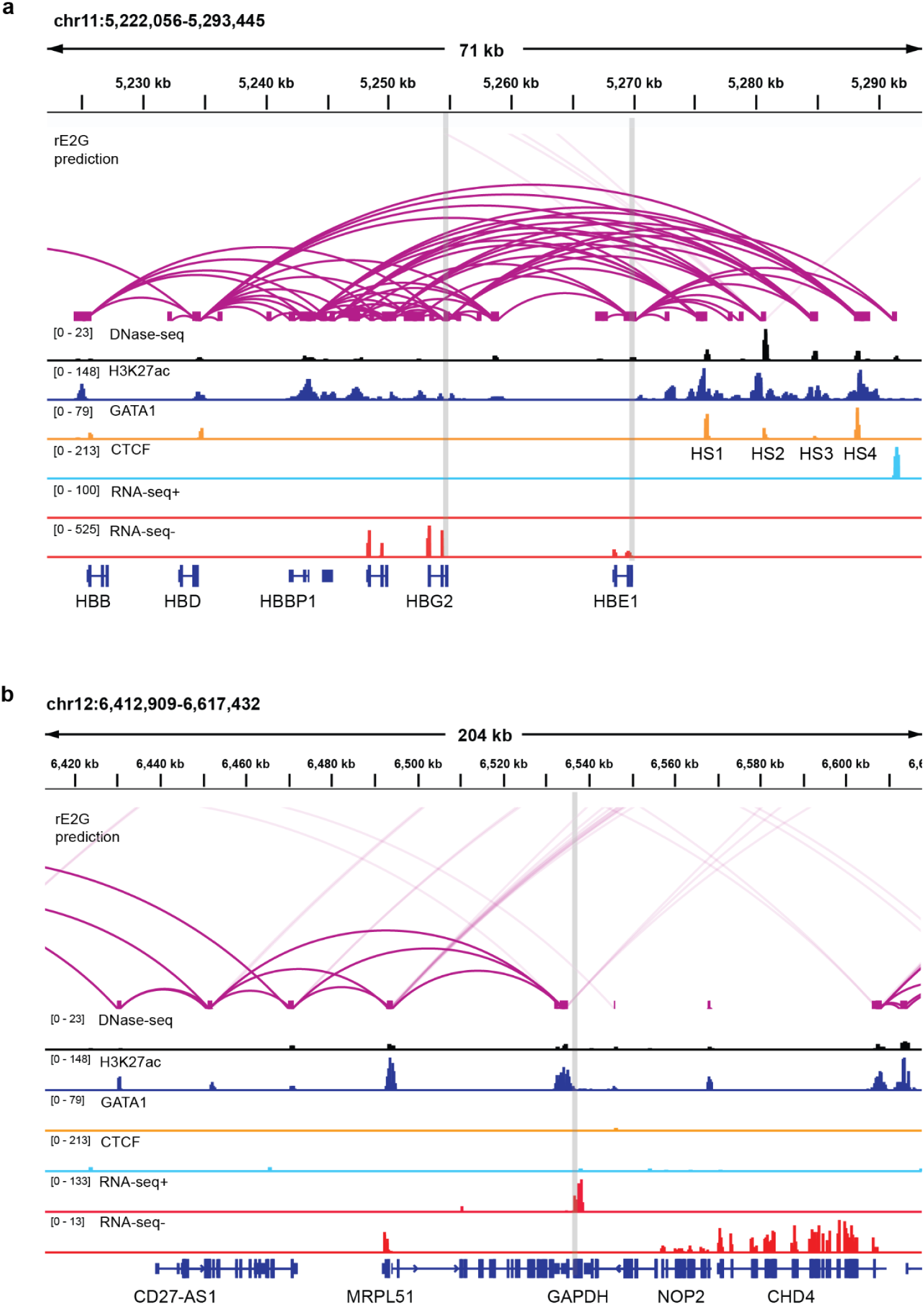
Genomic context of *HBG*, *HBE* and *GAPDH* promoters at their endogenous locus. **a-b**, Genomic context of *HBG*, *HBE* (a) and *GAPDH* promoter (b) at their endogenous locus in K562 cells. Tracks show: rE2G predicted E-P interactions, DNase-seq signal, ChIP-seq tracks for H3K27ac, GATA1 and CTCF, total RNA-seq data for plus/minus strand and RefSeq gene annotations. Promoter locations are highlighted with grey boxes. The four HSs at the LCR locus are labeled in (a).

**Extended Data Fig. 5:**
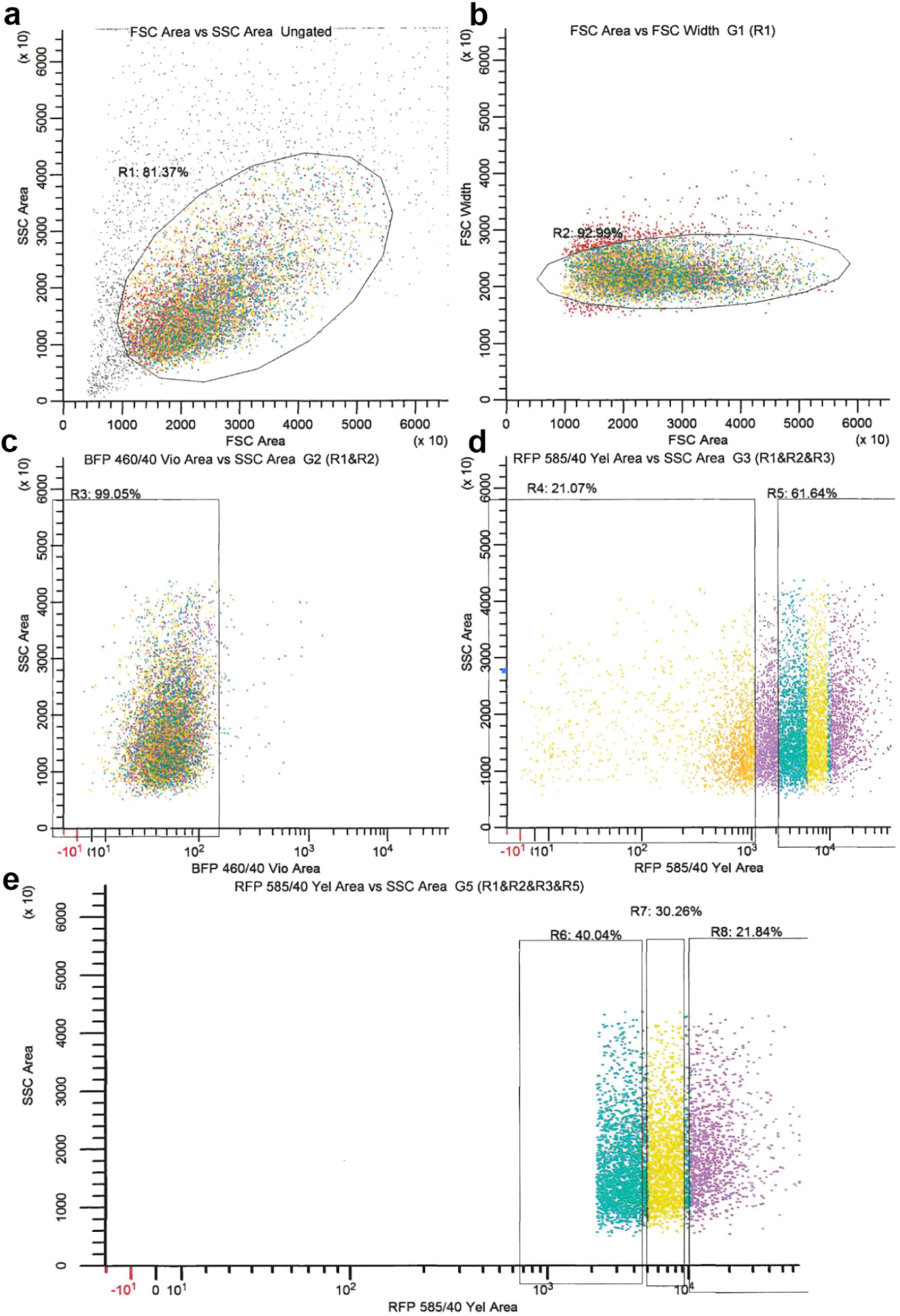
Example of the gating strategy (replicate 3 of the 6-distance LCR library). **a**, Gating for live cells based on Forward Scatter Area and Side Scatter Area. **b,** Gating for single cells based on Forward Scatter Area and Forward Scatter Width. **c,** Gating for BFP negative population to enrich cells with successful enhancer integration. **d,** Further gating for RFP to classify enhancer activity. The R4 gate is set as Bin1 for no/low activity enhancers, defined as those below the 75th percentile of the control cells with only basal promoter expression. R5 represents RFP positive population, defined as those above 90 percentile of the control cells with only basal promoter expression. **e,** R5 is further split into Bin2, Bin3 and Bin4, using a 40%:30%:20% distribution.

